# Deep Multimodal Graph-Based Network for Survival Prediction from Highly Multiplexed Images and Patient Variables

**DOI:** 10.1101/2022.07.19.500604

**Authors:** Xiaohang Fu, Ellis Patrick, Jean Y. H. Yang, David Dagan Feng, Jinman Kim

## Abstract

The spatial architecture of the tumour microenvironment and phenotypic heterogeneity of tumour cells have been shown to be associated with cancer prognosis and clinical outcomes, including survival. Recent advances in highly multiplexed imaging, including imaging mass cytometry (IMC), capture spatially resolved, high-dimensional maps that quantify dozens of disease-relevant biomarkers at single-cell resolution, that contain potential to inform patient-specific prognosis. However, existing automated methods for predicting survival typically do not leverage spatial phenotype information captured at the single-cell level, and current methods tend to focus on a single modality, such as patient variables (PVs). There is no end-to-end method designed to leverage the rich information in whole IMC images and all marker channels, and aggregate this information with PVs in a complementary manner to predict survival with enhanced accuracy. We introduce a deep multimodal graph-based network (DMGN) that integrates entire IMC images and multiple PVs for end-to-end survival prediction of breast cancer. We propose a multimodal graph-based module that considers relationships between spatial phenotype information in all image regions and all PVs, and scales each region–PV pair based on its relevance to survival. We propose another module to automatically generate embeddings specialised for each PV to enhance multimodal aggregation. We show that our modules are consistently effective at improving survival prediction performance using two public datasets, and that DMGN can be applied to an independent validation dataset across the same antigens but different antibody clones. Our DMGN outperformed state-of-the-art methods at survival prediction.

## 1. Introduction

The spatial organisation and structure of the tumour microenvironment surrounding tumour cells, and cellular phenotypic heterogeneity are associated with clinical outcomes and prognosis in cancer [1–3]. This information may be valuable for improving the accuracy of survival prediction models, which aim to predict the time to death events given an array of information regarding the patient. Recent techniques in highly multiplexed imaging mass cytometry (IMC) use isotope-tagged antibodies to capture maps of the spatial expression of over 30 biomarkers at the single-cell level [3–5]. These images contain detailed high-dimensional information and provide opportunities for further quantitative analysis, including survival prediction.

Traditional survival analysis typically relies on the Cox proportional hazards (CPH) model [6]. More recently, deep learning (DL) models have been successfully applied in a wide range of applications, including modelling the survival function. In contrast to conventional survival models, DL models do not require manual feature generation and selection, as they can automatically extract deep features from the inputs that are relevant to survival. As an example, DeepSurv [7] used a negative Cox partial likelihood loss to handle right-censored survival data. However, existing DL survival prediction methods typically focus on analysing a single input modality, especially patient variables (PVs), which include the patient’s clinical information such as age or gene expression [7–11]. These methods do not consider spatial phenotype information at the cellular level that is relevant to disease prognosis.

Multiplexed images have recently been leveraged to predict oestrogen receptor (ER) status [12]. This demonstrated the potential of a graph neural network (GNN) [13], which models objects as nodes and capture the relationships between them as edges, to represent cellular communities and cell-cell interactions. However, this approach is unimodal and does not consider PVs. It also relies on considerable image pre-processing to segment cells and create neighbourhoods, which adds extra labour and potential errors that would propagate to the survival analysis downstream.

### 1.1. Our contributions

- We introduce a deep multimodal graph-based network (DMGN) that integrates highly multiplexed IMC images and multiple PVs for survival prediction of cancer. To our knowledge, our method is the first to combine these two modalities for end-to-end survival prediction. The source code is available at https://github.com/xhelenfu/DMGN_Survival_Prediction.
- We propose a region-based multimodal relational module (RMRM) to combine features from both modalities. The RMRM graph considers relationships between all PVs and spatial phenotype information across the entire IMC image and all biomarkers. The relationship between each PV and spatial region is scaled based on its strength, which allows the RMRM to capture relationships in more detail and support explainability.
- We propose a patient variable embedding module (PVEM) to automatically generate embedded representations that are meaningful to each PV through auxiliary reconstruction tasks. These representations support the relational learning with image features in the RMRM and improve survival prediction.
- Our method does not require cell segmentation, marker selection, or any other input processing external to the basic DL data pipeline.

## 2. Methods

### 2.1. Datasets

We used the Molecular Taxonomy of Breast Cancer International Consortium (METABRIC) dataset [3, 4] as the primary dataset. IMC images and PVs were available for 456 patients (518 samples). The uncensored rate (observed death) was 45.2%, and the median survival time was 7.97 years. We used 9 variables as done by previous studies [7–10], comprising 5 clinical features (age at diagnosis, ER status, chemotherapy indicator (CT), hormone treatment indicator (HT), and radiotherapy indicator (RT)), and 4 gene indicators (MKI67, EGFR, PGR, and ERBB2). Invasive carcinoma cores of 0.6 mm were labelled with antibodies and quantified using a Hyperion Imaging Mass Cytometer (Fluidigm). The IMC images contained 50 channels with heights of 25-744 (average 464) pixels, and widths of 60-834 (average 478) pixels.

We used the Basel dataset [5] for further cross-validation and independent validation. IMC images and PVs were available for 285 patients (376 samples). The uncensored rate was 28.1%, and the median survival was 6.17 years. We used the same PVs as for the METABRIC dataset, except radiotherapy indicator (RT) was unavailable and was substituted by immune therapy (IT). Cores of 0.8 mm were analysed using a Hyperion Imaging System (Fluidigm). The IMC images contained 52 channels with heights of 223–883 (average 711) pixels, and widths of 178–1032 (average 746) pixels.

We used all censored and uncensored samples, and marker channels for both datasets in the model inputs. All images were resized to 176 × 176 pixels with bilinear interpolation. Details of the datasets are in Table 1.

**Table 1.** Details of the two public datasets used in this study.

|  | METABRIC | Basel |
| --- | --- | --- |
| Number of patient variables | 9 | 9 |
| Median censored time | 7.97 years | 6.17 years |
| Uncensored rate | 45.2% | 28.1% |
| Total number of patients | 456 | 285 |
| Total number of samples | 518 | 376 |
| Number of IMC channels | 50 | 52 |

### 2.2. Deep multimodal graph-based network (DMGN)

In our method, information from each modality is first extracted by two separate branches. IMC images are processed by a CNN backbone to output a grid of feature vectors that describe biomarker expressions in spatial regions (patches of pixels) across the image. PVs are processed by the PVEM to generate variable-specific embedding vectors. The feature vectors for both modalities are then integrated in the RMRM and used to predict survival. We present an overview of our method in Fig. 1.

**Fig. 1.**
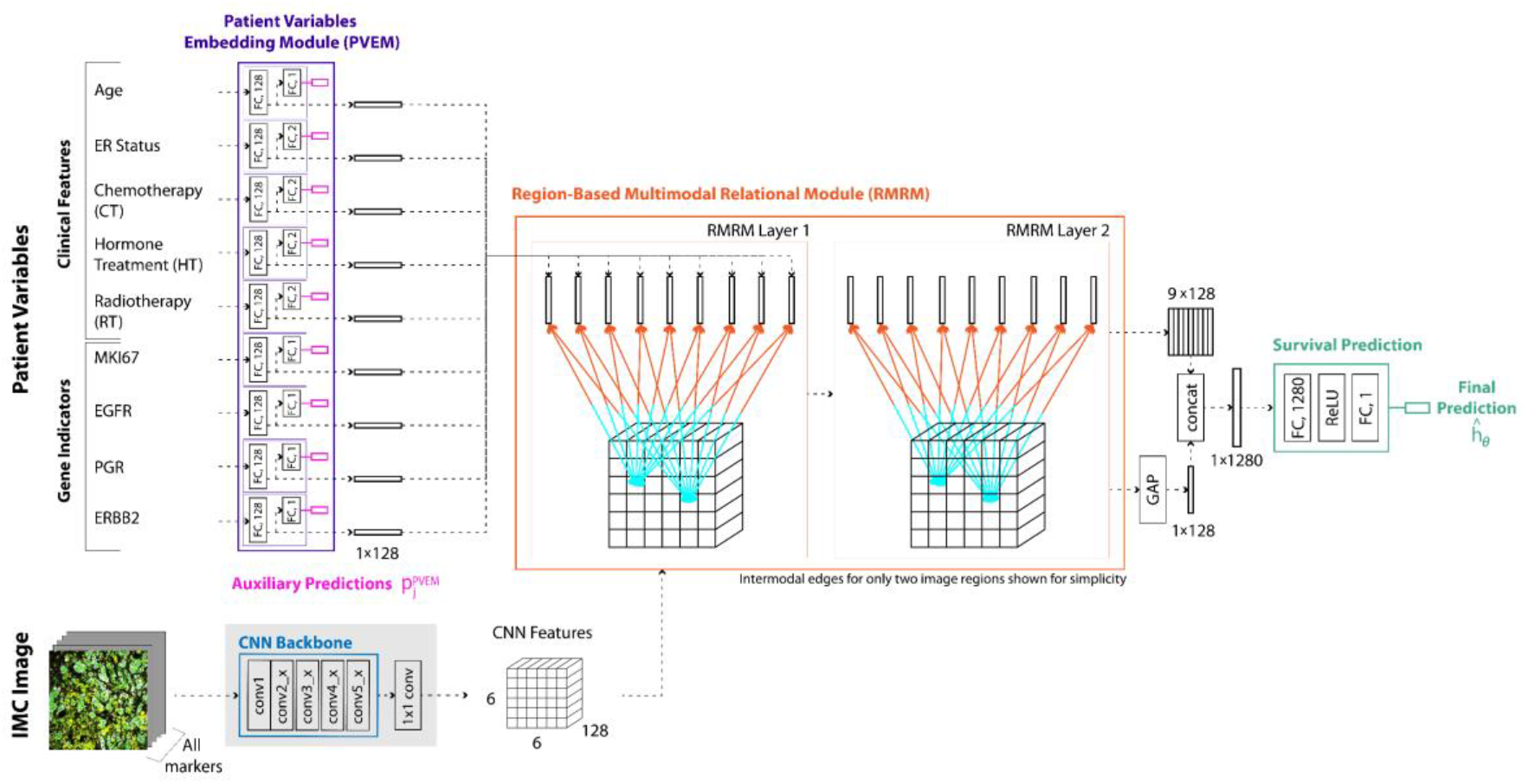
Proposed DMGN model for end-to-end survival prediction from PVs and IMC images. PVs are processed by the PVEM to produce embeddings. The IMC image is processed by a CNN backbone to create deep image features. Both modalities are combined in the RMRM and then used to predict survival.

We use the versatile ResNet-50 model [14] as the CNN backbone for visual feature extraction. We do not select for certain IMC markers; all channels in the IMC image are used. We insert a 1 × 1 convolution layer immediately after the ‘conv5_x’ block of the backbone to reduce the number of feature channels from 2,048 to 128. This minimises the risk of overfitting by reducing the number of parameters in the model. The output of the CNN backbone is a feature volume 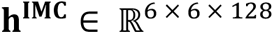 for each input image.

#### 2.2.1. Patient variable embedding module (PVEM)

Prior to multimodal relational learning, each PV is transformed into feature embeddings by the PVEM. There are 9 units in the module, each specialised for a PV. The units consist of fully connected (FC) layers arranged to perform autoencoding of the input variable, i.e., they learn to reconstruct the input variable.

We define the units to output PV embeddings and reconstructions as:

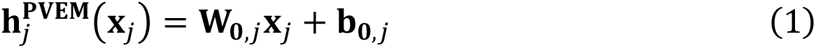

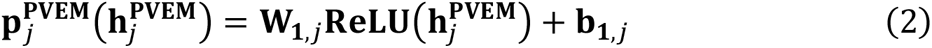

where 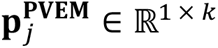 denotes the prediction for variable *j*. 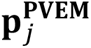 is a single scalar *(k* = 1) for continuous PVs (age, MKI67, EGFR, PGR, and ERBB2), or class probabilities (*k* = 2) for categorical variables (ER status, CT, HT, and RT/IT) with softmax activation. 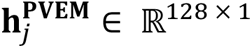 is the variable-specific feature vector embedding. The PVEM unit of a category is characterised by a set of trainable weights and biases of two FC layers, comprising 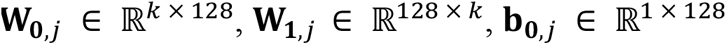, and 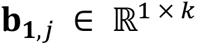.

During training, **p^PVEM^** is used to compute auxiliary reconstruction losses *L^PVEM^* that contribute to the total loss. *L^PVEM^* is the sum of the mean squared errors of the continuous variables and cross-entropy losses of the categorical variables.

By minimising the auxiliary losses, PVEM ensures that **h^PVEM^** is a meaningful representation of each PV. This assists with multimodal relational processing in the RMRM. The auxiliary predictions are only used during model training and have no role at inference time.

#### 2.2.2. Region-based multimodal relational module (RMRM)

The RMRM structure considers relationships between the spatial phenotype information in each IMC image region and all 9 PVs. This structure is designed to encode all pairs of relationships between each IMC region and PV, and allows each pair to be scaled based on its relevance to survival. We use two graph convolutional network (GCN) layers with the same graph topologies to integrate features from both modalities in the RMRM. We represent each spatial region as separate nodes in the graph, and each PV is further represented by individual nodes. The input feature vector for the node of IMC region *i* is 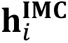, and PV *j* is 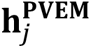. There are 45 nodes in each RMRM layer (36 IMC regions and 9 PVs).

The encoded features of the nodes are transformed and passed between the two modalities via bidirectional edges and graph convolutions. In this way, IMC features are integrated with PV features, and PV features are also integrated with IMC features. Graph convolutions allow each node to receive the aggregated feature vectors of its neighbouring (connected) nodes. Each IMC region receives the features of all PVs (through PV–IMC edges), and each PV receives the features of all IMC regions (through IMC–PV edges). Additionally, there is a self-edge for each node which preserves some features of each own node during a graph convolution. The adjacency matrix that describes the RMRM is presented in Fig. 2.

**Fig. 2.**
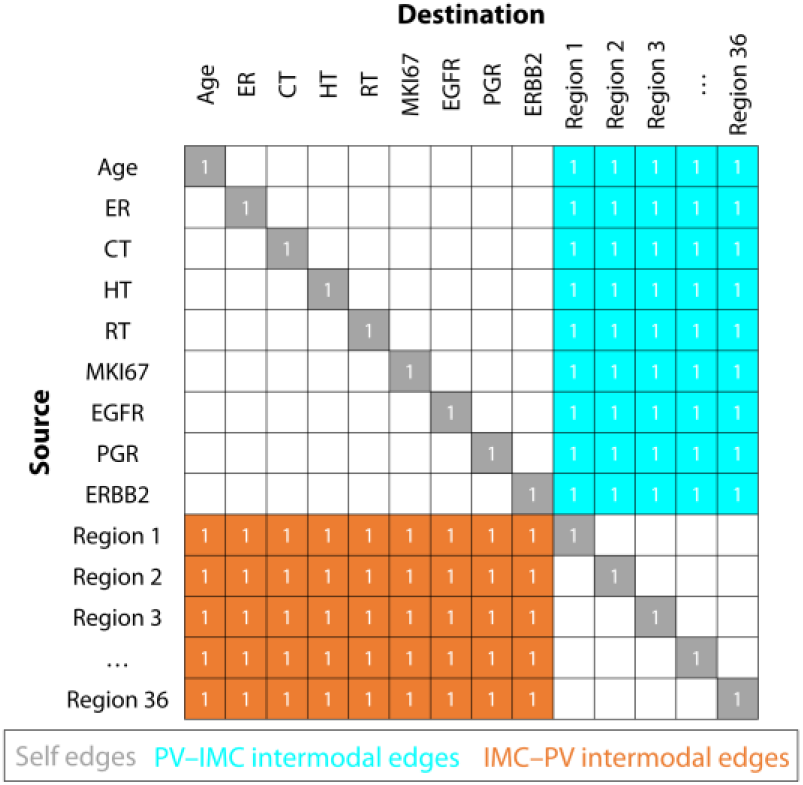
RMRM adjacency matrix. Filled squares are edges between source and destination nodes. Blank squares represent the absence of an edge (zeros are omitted for simplicity). All 36 image regions are connected to the 9 PVs by bidirectional edges.

Each graph convolution in the RMRM linearly transforms the neighbouring feature vectors via learned weights, then sums the resulting vectors and applies an activation function to the output. In a standard GCN, the neighbouring nodes are weighted equally towards the central node. We used the graph attention network (GAT) [15] approach to allow the contributions of each neighbour node towards the central node to be adaptive, rather than equal and fixed. This allows the most important information to be prioritised. GAT operates on the graph layers of the GRM with topologies and connections as described in Fig. 2. The graph attentional layer computes normalised attention coefficients via a self-attention mechanism:

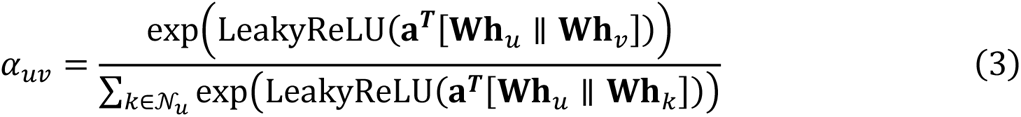

where 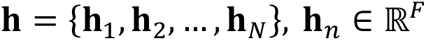 represents the input feature vectors to the GAT layer with *N* = 45, corresponding to the 9 PVs and 36 IMC regions (i.e., 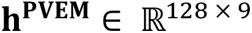 and 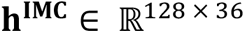), *a_uv_* is the attention coefficient for node *v* to *u,* and *N_u_* denotes the neighbourhood nodes of *u*. The feature vectors are linearly transformed via 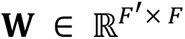, the weight matrix shared for every node, then by a 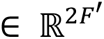, the weight vector of a single-layer feedforward neural network of the attention mechanism, with || representing concatenation. LeakyReLU nonlinearity (with a slope of 0.2 for inputs < 0) followed by softmax normalisation are then applied.

The updated feature vectors, 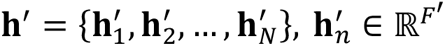, can then be computed by proportionally aggregating over neighbouring nodes:

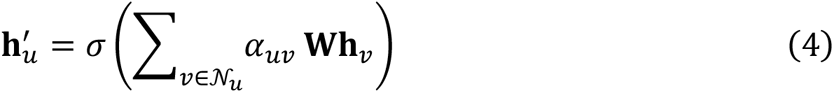

where σ is the exponential linear unit (ELU) nonlinearity [16].

Furthermore, we used the multi-head attention mechanism for better stability, computing multiple independent attention coefficients for each node that are aggregated via concatenation:

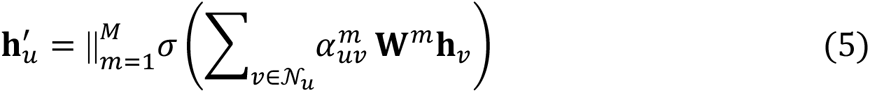

where, for the *m*-th attention head, 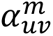 are the normalised attention coefficients and **W***^m^* are the weights of the input linear transformation. In the first layer, we use 8 attention heads per node and F_0_ = 8, thereby computing a total of 64 features, as done in the original paper [15]. For the second layer, we compute F_1_ = 128 features with a single attention head.

#### 2.2.3. Survival prediction and training

The RMRM outputs of the IMC feature volumes 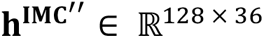 are subsequently vectorised by a global average pooling (GAP) layer [17], producing a vector of 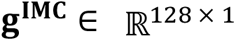 per image. This vector is then concatenated with all RMRM PV output vectors, resulting in one vector with dimensionality 1,280 × 1 per input sample. The last group of layers comprises an FC layer, ReLU activation, and a final output FC layer with linear activation.

The output of the model is a scalar 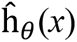(*x*) that estimates the log-risk function in the Cox model [7]. The model is trained by minimising the sum of 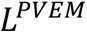 and mean negative Cox partial likelihood loss 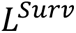, which is suitable for right-censored survival data:

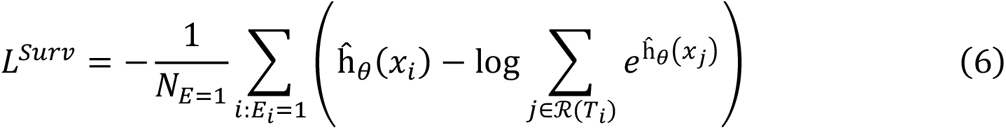

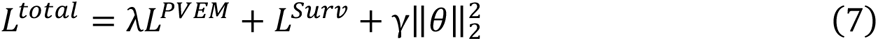

where *x_i_* denotes the input IMC image and PVs of sample *i, θ* represents the model parameters, *E* indicates an observed event (*N*_*E*=1_ is the number of patients with recorded death), *T* is the overall survival time (for *E* = 1) or censored time (for *E* = 0), and 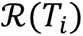 are the patients with *T* ≥ *T_i_*. λ is a hyperparameter to scale *L^PVEM^*, and *γ* is the L_2_ regularisation weight.

### 2.3. Implementation details

The following implementation and hyperparameter choices were kept consistent for all experiments to ensure a fair comparison. The networks were trained end-to-end from scratch for 300 epochs with a batch size of 8. We employed the Adam optimiser [18] to minimise the training loss (Eq. 7) at a fixed learning rate of 0.0001, with a first moment estimate of 0.9 and second moment estimate of 0.999. We set λ to 4, as determined empirically (evaluation in Supplementary Table S1), and γ to 0.0001. Weights of the convolutional layers were initialised using He et al.’s method [19], while biases were initialised to zero. Glorot (Xavier) initialisation [20] was used for FC layers. Dropout was not used in any experiment.

The channels of the images were mean-subtracted and normalised to unit variance using the training set means and standard deviations. We employed standard online (on- the-fly) image data augmentation by randomly applying a flip (horizontal or vertical), or rotation (of 90, 180 or 270 degrees) to the input images. The order of training examples was shuffled for every epoch. All networks were implemented using the PyTorch framework [21]. Both training and testing were performed with a 12GB NVIDIA GTX Titan V GPU.

### 2.4. Evaluation metrics

Survival prediction performance of the methods was evaluated using Harrell’s concordance index (C-index) [22], which is one of the standard evaluation metrics in survival analysis. The C-index ranges from 0 to 1.0, where 1.0 indicates perfect ranking of predicted death times, and 0.5 indicates random predictions.

The log-rank test was used to assess the difference between two Kaplan-Meier survival curves. A log-rank *p*-value below 0.05 is considered statistically significant.

### 2.5. Ablation study and independent validation

We performed an ablation study to validate each component of our DMGN by assessing their effects on survival prediction performance. We first determined the performance using only a single modality—either IMC images or PVs. The model for image-only inputs consisted of a ResNet-50 backbone with the same survival prediction output layers as in the DMGN. The model for the PVs consisted of the PVEM (but without auxiliary reconstruction losses) and the same survival prediction output layers.

We then benchmarked conventional approaches to combine multimodal information, including concatenation of multimodal feature vectors, and attention-weighted aggregation. Attention is a common approach in many multimodal applications that use weights to select and combine relevant features from both modalities [23–25].

We assessed the performance of our DMGN without auxiliary PVEM reconstruction losses. Furthermore, we investigated a modified RMRM that operates on 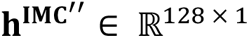 instead of 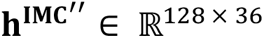, by moving GAP to just before the RMRM instead of just after. This effectively removes all spatial information in the RMRM by compressing the image features to a single feature vector.

We performed 5-fold cross-validation on both datasets to show the effectiveness of our method and that it outperforms previous methods using in-silico measurements in terms of higher C-index. We randomly assigned 80% of the patients for training and the remaining 20% for testing in each fold. We also carried out independent validation where the model was trained using METABRIC data but validated using all available Basel data. We used markers common to both datasets and removed RT from METABRIC data and IT from Basel data. There was no additional input pre-processing or model finetuning for this independent validation.

### 2.6. Comparison to existing methods

We benchmarked our proposed DMGN to existing survival prediction methods using both datasets. State-of-the-art methods for survival prediction on the METABRIC dataset included DeepHit [11], DeepSurv [7], BroadSurv [9], DAGSurv [10], and AttentionSurv [8]. We report the C-index for the 3-year predicted survival for DAGSurv as it is a discrete-time survival model, following previous survival analysis studies [26, 27]. All these methods were designed for single modality inputs and use the same 9 PVs as our DMGN (age, ER status, CT, HT, RT, MKI67, EGFR, PGR, and ERBB2), except for DeepHit, which used 21 variables. We also report results for the CPH model [6] and random survival forest (RSF) [28].

We used the CPH model [6] with various input modality combinations and regularisation as the benchmark for the Basel dataset, due to a lack of recent methods. We used either L_1_, L_2_, or L_1_+L_2_ regularisation strategies. The inputs were either cell type proportions, PVs, or both cell type proportions and PVs. Cell type proportions of each IMC image were computed using metacluster cell type counts [5], divided by the total cell count for the image.

## 3 Results

### 3.1. Ablation study and validation

The prediction performance of our DMGN was consistently superior compared to the various DL baselines for the two datasets (Table 2). The single modality (image or PVs only) results indicate that PVs contain more useful information for survival prediction compared to images. The higher scores of all multimodal approaches compared to single modality approaches demonstrate the importance in using both modalities.

**Table 2.** Prediction performance (C-index) of the proposed DMGN compared to DL baselines with different inputs and fusion methods. Bold indicates the best performance for each dataset.

| Method | METABRIC | Basel |
| --- | --- | --- |
| IMC image only | 0.6581 | 0.6769 |
| PVs only | 0.7147 | 0.7078 |
| Multimodal, concatenation | 0.7174 | 0.7131 |
| Multimodal, attention | 0.7327 | 0.7242 |
| Multimodal, DMGN (no aux losses) | 0.7398 | 0.7354 |
| Multimodal, DMGN (no spatial) | 0.7399 | 0.7359 |
| Multimodal, DMGN (proposed) | **0.7484** | **0.7479** |

In both datasets the concatenation approach improved performance slightly compared to using only PVs. Concatenation is a suboptimal approach to combine multimodal information as it does not extract any further useful relationships from the two modalities that would be more informative compared to single modality information. The improvements from attention-weighted aggregation were more substantial, as this approach prioritises the multimodal features according to their importance for survival prediction.

All DMGN variants were superior to concatenation and attention-based aggregation. Performance of the DMGN was lower without auxiliary reconstruction losses. This finding supports the motivation and design of reconstruction losses, i.e., they allow the PVEM output features to be more descriptive of each PV, thereby allowing more meaningful relational learning with image features and ultimately aiding survival prediction.

DMGN performance was also inferior without spatial image information during relational processing in the RMRM. Our proposed topology considers different relationships between each PV to various regions of the image. This allows multimodal relationships to be extracted in more detail. The ablation study shows that the spatial simplification of IMC images has a deleterious effect in the present multimodal fusion task.

To further illustrate the utility of our model, the Kaplan-Meier survival curves for one testing fold from the METABRIC (left) and Basel (right) datasets are shown in Fig. 3. Our DMGN successfully stratified the patients into two significantly different high and low risk groups (log-rank *p* = 3 ×10^−21^ for METABRIC, *p* = 2×10^−9^ for Basel). The curves for the other folds displayed similarly significant separation between the two groups (all log-rank *p* < 10^−5^).

**Fig. 3.**
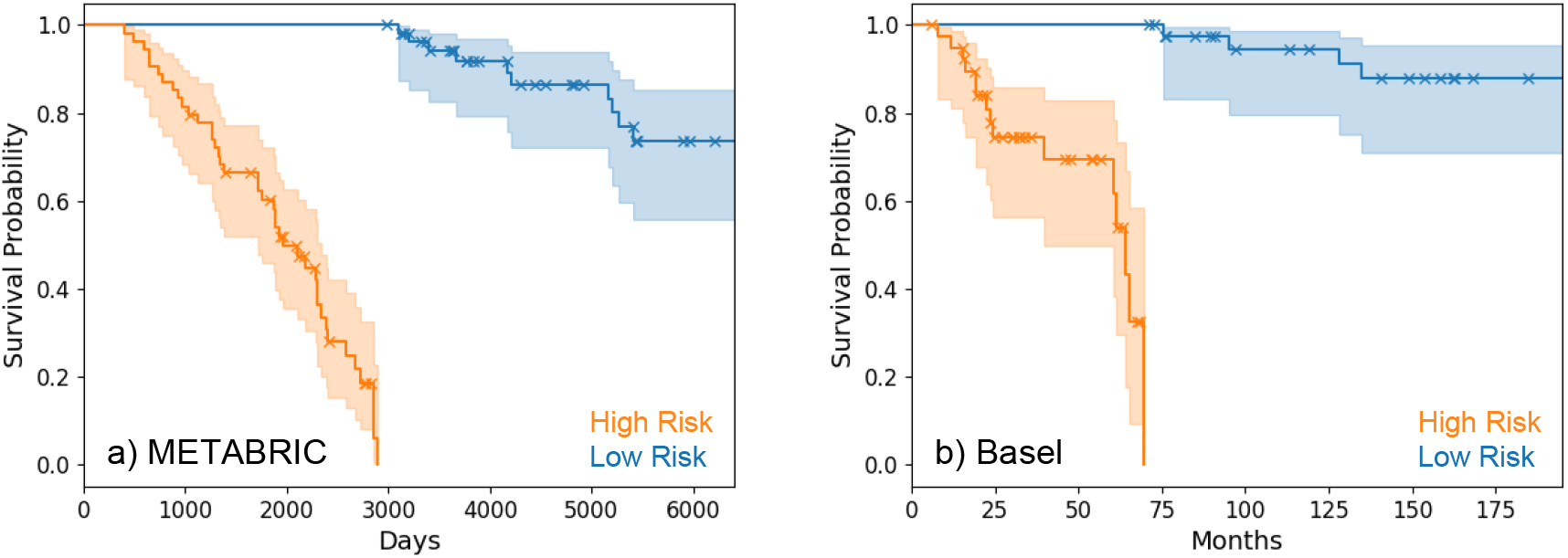
Kaplan-Meier survival curves. DMGN predictions for high risk (higher than or equal to median) and low risk (lower than median) groups for a) METABRIC and b) Basel datasets for one testing fold. ‘ × ‘ indicates a censored patient. Predicted KM curves are significantly different for high and low risk groups (log-rank *p* = 3×10^−21^ for METABRIC, *p* = 2×10^-9^ for Basel).

The independent validation performance for our DMGN that was trained on METABRIC data but tested on Basel data was lower than our model that was both trained and tested on Basel data (Table 3). This finding is not surprising for DL, since learned weights adapt to the characteristics of the training set and typically do not generalise to new data at the same accuracy [29]. The reduction in independent validation performance would also be compounded by heterogeneous characteristics between the datasets, and differences in panel design and imaging protocols [3–5].

**Table 3.** Validation of the proposed DMGN on an independent test set.

| Method | C-index |
| --- | --- |
| Test METABRIC (trained METABRIC) | 0.7380 |
| Test Basel (trained Basel) | 0.7146 |
| Independent test Basel (trained METABRIC) | 0.6312 |

### 3.2. Comparison to existing methods

Our DMGN had superior survival prediction performance compared to all existing methods for both datasets (Table 4 and Table 5). Performance of CPH with different input modalities on the Basel dataset (Table 5) followed the same pattern as the DL ablations experiments (Table 2), where image-only inputs resulted in the lowest performance. Performance using PV-only inputs was higher but was surpassed by multimodal inputs.

**Table 4.** Comparison of the proposed DMGN to existing methods on the METABRIC dataset.

| Method | C-index |
| --- | --- |
| CPH | 0.632 |
| RSF | 0.620 |
| DeepSurv [7] | 0.654 |
| DeepHit [11] | 0.691 |
| BroadSurv [9] | 0.643 |
| DAGSurv [10] | 0.689 |
| AttentionSurv [8] | 0.708 |
| DMGN (proposed) | **0.748** |

**Table 5.** Comparison of the proposed DMGN to existing methods on the Basel dataset.

| Method | C-index |
| --- | --- |
| CPH (cell type proportions, L<sub>2</sub>) | 0.584 |
| CPH (PVs, L<sub>2</sub>) | 0.686 |
| CPH (cell type proportions + PVs, L<sub>1</sub>) | 0.696 |
| CPH (cell type proportions + PVs, L<sub>2</sub>) | 0.698 |
| CPH (cell type proportions + PVs, L<sub>1</sub> + L<sub>2</sub>) | 0.697 |
| DMGN (proposed) | **0.748** |

### 3.3. Weights of patient variables

We analysed the weights of the RMRM connections between PVs and image regions (PV–IMC intermodal edges, cyan in Fig. 2). The average weights of the test samples in all cross-validation folds and their rankings for both datasets are presented in Table 6. The weights revealed different patterns and characteristics of the two cohorts. The rankings for the two cohorts were different, e.g., the top two PVs were ‘age’ and ‘MKI67’ for METABRIC but were ‘ERBB2’ and ‘HT’ for Basel. The differences in the weights of the PVs were smaller for Basel compared to METABRIC.

**Table 6.**
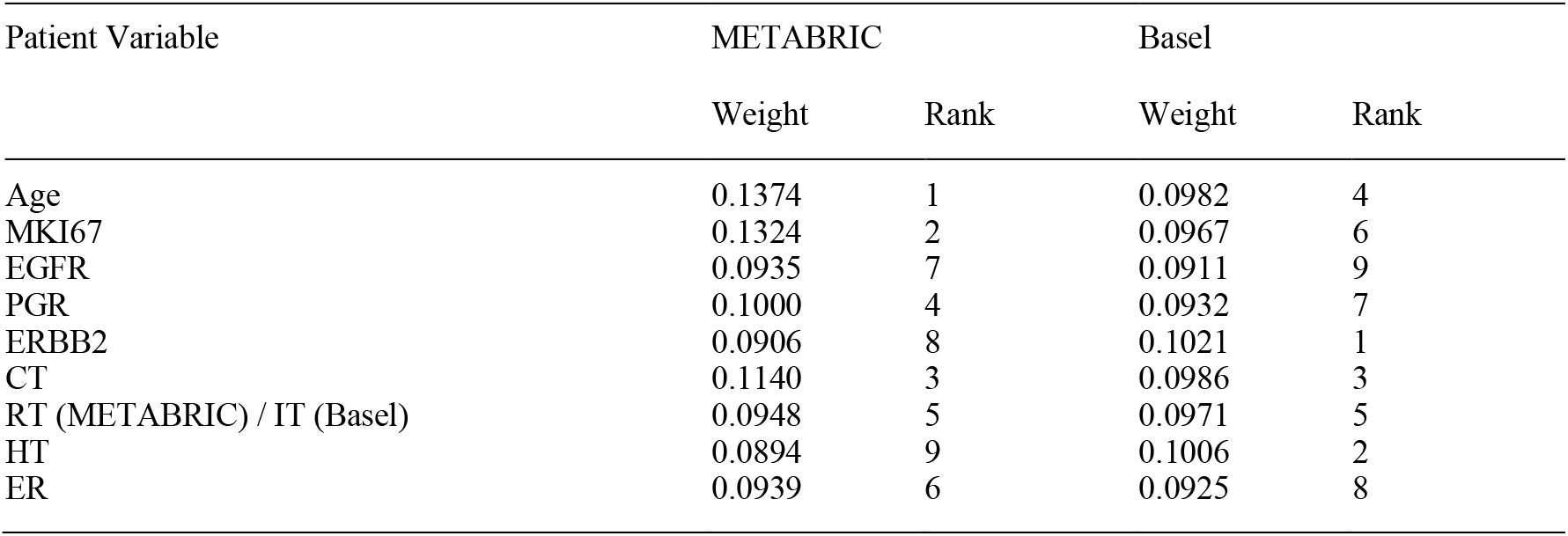
Average RMRM weights of the PVs to image regions and their rank in descending order.

| Patient Variable | METABRIC |  | Basel |  |
| --- | --- | --- | --- | --- |
|  | Weight | Rank | Weight | Rank |
| Age | 0.1374 | 1 | 0.0982 | 4 |
| MKI67 | 0.1324 | 2 | 0.0967 | 6 |
| EGFR | 0.0935 | 7 | 0.0911 | 9 |
| PGR | 0.1000 | 4 | 0.0932 | 7 |
| ERBB2 | 0.0906 | 8 | 0.1021 | 1 |
| CT | 0.1140 | 3 | 0.0986 | 3 |
| RT (METABRIC) / IT (Basel) | 0.0948 | 5 | 0.0971 | 5 |
| HT | 0.0894 | 9 | 0.1006 | 2 |
| ER | 0.0939 | 6 | 0.0925 | 8 |

## 4 Discussion

### 4.1. Ablation study and validation

Our modules provide advantages that are not feasible in conventional multimodal fusion approaches such as with concatenation and attention. The RMRM captures multimodal relationships across different spatial regions of the image, and scales the contributions from each feature to prioritise the most important relationships. Multimodal aggregation is also carried out with meaningful representations of the PVs due to the PVEM. The ablation study (Table 2) validated the benefits of using both modalities (IMC images and PVs), and the effectiveness of our DMGN architecture with PVEM and RMRM for survival prediction.

Our DMGN has demonstrated competence in discriminating high- and low-risk groups (Fig. 3), which is practically beneficial as this knowledge may be used to guide personalised patient management. However, the performance of DMGN may be limited when directly applied to an independent cohort with different data characteristics (Table 3). While it was encouraging that our DMGN was robust to a reordering and omission of markers when trained and tested on IMC experiments with similar but ultimately different panel designs, standardised imaging protocols or pre-processing would clearly help to ensure comparable image intensity distributions and alleviate performance reductions [30].

### 4.2. Comparison to existing methods

In addition to the contributions provided by our PVEM and RMRM, our DMGN is distinct from existing methods in our use of IMC images in addition to PVs. These images spatially quantify multiple clinically established breast cancer biomarkers such as MKI67 and HER2, and characterise spatial cellular heterogeneity, composition, and organisation [3–5], which are potential indicators of cancer prognosis [1, 2]. The use of a single modality in survival analysis (i.e., PVs) as done by previous studies is therefore limited, and our results show that the value in multiplexed images should be exploited. Our model can leverage both modalities, and aggregate PVs and spatial features across the image to consistently outperform the state-of-the-art (Table 4 and Table 5).

### 4.3. Weights of patient variables

The PVEM allows a level of explainability as the relative strengths between the PVs and image regions can be assessed according to the weights of each connection in the graph. This ability of the PVEM may be used to understand the contributing factors towards survival predictions and signify samples for further analysis. For example, image regions with particularly high or low PVEM weights to certain PVs may be quantitatively analysed at the cellular level.

### 4.4. Future work

While our DMGN was specifically optimised and validated on IMC data, there is potential for it to be extended and applied to other multiplexed imaging assays [31]. This could include other proteomic assays such as co-detection by indexing (CODEX) [32] and cyclic immunofluorescence (CycIF) [33] which can measure a similar number of protein markers to IMC but can measure larger regions of tissue. It could also include transcriptomic assays such as sequential Fluorescence in situ Hybridization (seqFISH+) [34] and high-definition spatial transcriptomics [35] that can measure an order of magnitude more features.

PV embeddings may be integrated with image features at different stages in the CNN. Such a design may be more effective at leveraging image features at different scales. Future iterations of the DMGN can include components designed to increase the explainability and interpretability of the architecture. Cell segmentation masks, or cell segmentation as a secondary objective may be incorporated into the architecture to delineate relationships between IMC markers and PVs that are inside or outside cell areas. This may be beneficial for performance or explainability.

## 5. Conclusion

We introduced an end-to-end DL model for survival prediction that integrates multiple PVs with biomarker expressions from highly multiplexed images. Our RMRM aggregates features from all regions and channels of IMC images with PVs, and scales each intermodal relationship to improve survival prediction performance. Our PVEM can automatically produce embedded representations for each PV to support intermodal learning and survival prediction. Our method showed potential to be directly applied to an independent dataset without further training, and consistently surpassed existing methods for predicting the survival of breast cancer patients.

## Acknowledgements

The authors thank their colleagues at The University of Sydney and the Sydney Precision Data Science Centre for their support and intellectual engagement.

## Data availability

All data used in this study are publicly available. The METABRIC dataset [3, 4] may be obtained from https://idr.openmicroscopy.org/ (accession code idr0076). The Basel dataset [5] may be obtained from https://doi.org/10.5281/zenodo.3518284.

## Supplementary Materials

**Table S1.**
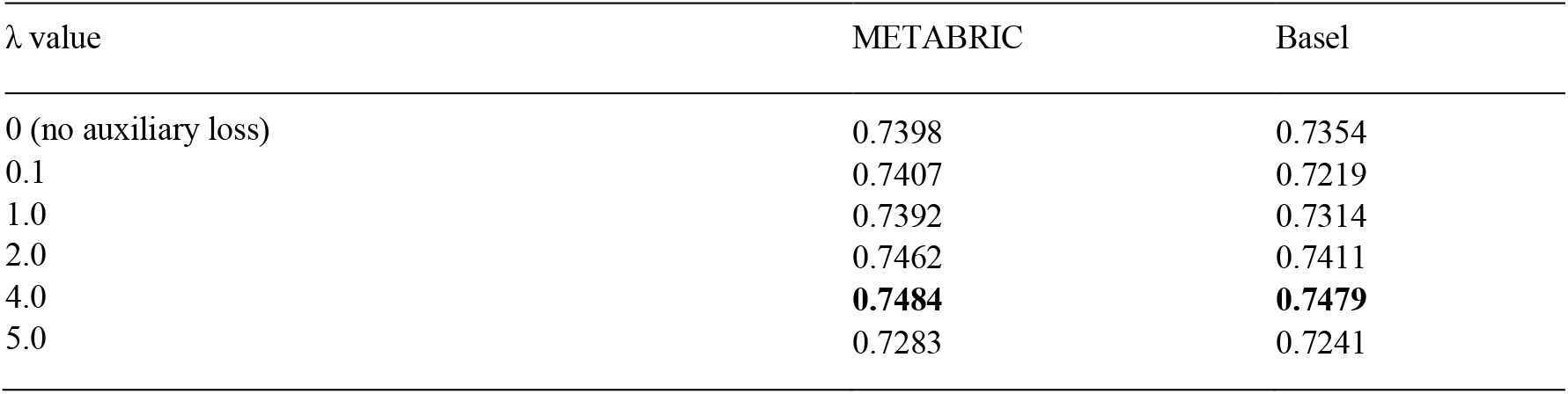
Prediction performance (C-index) of the proposed DMGN with various λ.

