## Supplementary Table S1 for "Deep Multimodal Graph-Based Network for Survival Prediction from Highly Multiplexed Images and Patient Variables"

**Table S1.** Prediction performance (C-index) of the proposed DMGN with various λ.

| λ value | METABRIC | Basel |
|---|---|---|
| 0 (no auxiliary loss) | 0.7398 | 0.7354 |
| 0.1 | 0.7407 | 0.7219 |
| 1.0 | 0.7392 | 0.7314 |
| 2.0 | 0.7462 | 0.7411 |
| 4.0 | **0.7484** | **0.7479** |
| 5.0 | 0.7283 | 0.7241 |
